# Retrograde suppression of post-tetanic potentiation at the mossy fiber-CA3 pyramidal cell synapse

**DOI:** 10.1101/2020.10.23.352377

**Authors:** Sachin Makani, Stefano Lutzu, Pablo J. Lituma, David L. Hunt, Pablo E. Castillo

**Affiliations:** Dominick P. Purpura Department of Neuroscience, Albert Einstein College of Medicine, Bronx, NY 10461; Department of Psychiatry and Behavioral Sciences, Albert Einstein College of Medicine, Bronx, NY 10461.; Current address: Center for Neural Science and Medicine, Cedars-Sinai Medical Center, Los Angeles, CA 90048

## Abstract

In the hippocampus, the excitatory synapse between dentate granule cell axons – or mossy fibers (MF) – and CA3 pyramidal cells (MF-CA3) expresses robust forms of short-term plasticity, such as frequency facilitation and post-tetanic potentiation (PTP). These forms of plasticity are due to increases in neurotransmitter release, and can be engaged when dentate granule cells fire in bursts (e.g. during exploratory behaviors) and bring CA3 pyramidal neurons above threshold. While frequency facilitation at this synapse is limited by endogenous activation of presynaptic metabotropic glutamate receptors, whether MF-PTP can be regulated in an activity-dependent manner is unknown. Here, using physiologically relevant patterns of mossy fiber stimulation in acute mouse hippocampal slices, we found that disrupting postsynaptic Ca^2+^ dynamics increases MF-PTP, strongly suggesting a form of Ca^2+^-dependent retrograde suppression of this form of plasticity. PTP suppression requires a few seconds of MF bursting activity and Ca^2+^ release from internal stores. Our findings raise the possibility that the powerful MF-CA3 synapse can negatively regulate its own strength not only during PTP-inducing activity typical of normal exploratory behaviors, but also during epileptic activity.

**SIGNIFICANCE STATEMENT:** The powerful mossy fiber-CA3 synapse exhibits strong forms of plasticity that are engaged during location-specific exploration, when dentate granule cells fire in bursts. While this synapse is well-known for its presynaptically-expressed LTP and LTD, much less is known about the robust changes that occur on a shorter time scale. How such short-term plasticity is regulated, in particular, remains poorly understood. Unexpectedly, an *in vivo*-like pattern of presynaptic activity induced robust post-tetanic potentiation (PTP) only when the postsynaptic cell was loaded with a high concentration of Ca^2+^ buffer, indicating a form of Ca^2+^–dependent retrograde suppression of PTP. Such suppression may have profound implications for how environmental cues are encoded into neural assemblies, and for limiting network hyperexcitability during seizures.

## INTRODUCTION

Mossy fibers (MFs), the axonal projections of dentate granule cells (GC), provide a strong excitatory input onto hippocampal CA3 pyramidal neurons (Henze et al., 2000; Nicoll and Schmitz, 2005). The MF-CA3 synapse is well known for exhibiting uniquely strong forms of short-term potentiation, including paired-pulse facilitation (PPF) and frequency facilitation, which last milliseconds to seconds (Salin et al., 1996). More intense periods of high-frequency stimulation typically elicit post-tetanic potentiation (e.g. MF-PTP), which decays over several minutes (Griffith, 1990). These forms of plasticity are commonly attributed to an increase in presynaptic release probability (Pr) (Zucker and Regehr, 2002; Nicoll and Schmitz, 2005; Regehr, 2012), a change in the readily releasable pool (Vandael et al., 2020), and transiently convert the synapse from a high- to lower-pass filter (Abbott and Regehr, 2004). In behaving rodents, during place field activation, GCs can fire in high-frequency bursts (Pernia-Andrade and Jonas, 2014; Diamantaki et al., 2016; GoodSmith et al., 2017; Senzai and Buzsaki, 2017), driving CA3 pyramidal neurons above threshold (Henze et al., 2002). Thus, frequency facilitation and MF-PTP could have a profound impact on how memory traces are encoded into CA3 neural ensembles.

Given the apparent ease with which these robust forms of presynaptic potentiation are elicited at the MF-CA3 synapse, one might expect a process by which the connection is negatively regulated. In fact, there is evidence that endogenously released glutamate transiently suppresses frequency facilitation via presynaptic group II metabotropic glutamate receptors (II-mGluRs) (Scanziani et al., 1997; Toth et al., 2000; Kwon and Castillo, 2008a). It is unknown, however, whether forms of longer-lasting plasticity, such as MF-PTP, are also curtailed in an activity-dependent manner.

In the present study we performed whole-cell recordings from CA3 pyramidal neurons in mouse hippocampal slices and mimicked physiologically-relevant activity patterns by stimulating GCs with a brief, high-frequency bursting paradigm. To our surprise, MF-PTP was readily observed when the postsynaptic cell was dialyzed with a solution containing high Ca^2+^ buffering properties, but was nearly absent when the recording solution had a lower, more physiological Ca^2+^ buffering capacity. These findings strongly suggest that MF-PTP is normally suppressed by a Ca^2+^-dependent retrograde mechanism. Such negative feedback could not only enable homeostatic regulation of MF-CA3 synaptic strength, ensuring that the essential filtering properties of the connection are maintained during ongoing activity, but also prevent runaway network excitability, as may occur during epileptic activity.

## METHODS

### Slice Preparation

Animal procedures were approved by our Institutional Animal Care and Use Committee and adhered to National Institutes of Health guidelines. C57BL mice (18-28 days old) were deeply anesthetized with isoflurane, decapitated, and brains rapidly removed and both hippocampi were dissected. Transverse hippocampal slices (400 μm) were cut on a DTK-2000 microslicer (Dosaka, Kyoto, Japan) or a Leica VT1200 S vibratome, in ice-cold cutting solution containing (in mM): 215 sucrose, 2.5 KCl, 20 glucose, 26 NaHCO_3_, 1.6 NaH_2_PO_4_, 1 CaCl_2_, 4 MgSO_4_, and 4 MgCl_2_. After 10 minutes of incubation at room temperature, the cutting solution was exchanged for the artificial cerebrospinal fluid (ACSF) containing 124 NaCl, 2.5 KCl, 10 glucose, 26 NaHCO_3_, 1 NaH_2_PO_4_, 2.5 CaCl_2_, and 1.3 MgSO_4_. Both cutting and ACSF solutions were saturated with 95% O_2_ and 5% CO_2_ (pH 7.4). The slices recovered at room temperature for at least 1.5 hour before recording.

### Electrophysiology

Slices were transferred to a recording chamber and perfused with ACSF (2 ml/min). Recordings were done at 25°C, except for the experiments included in **Fig 1a, 6e–h,** and those that used BoTX that were performed at 32°C. The recording pipette was filled with an internal solution containing (in mM): 112 K-gluconate, 17 KCl, 0.04 CaCl_2_, 10 HEPES, 2 Mg-ATP, 0.2 GTP, 10 NaCl, and 0.1 EGTA (pH 7.2) (290–295 mOsm). For the experiments using 10 mM EGTA or BAPTA, the concentration of K-gluconate was reduced to compensate for osmolarity. The pipette resistance ranged from 3-4 MΩ. Series resistance (6–15 MΩ) was monitored throughout the experiment, and those experiments in which the series resistance changed by more than 10% were not included for analysis. Patch pipettes were pulled on a PP-830 vertical puller (Narishige, Tokyo, Japan).

**Figure 1.**
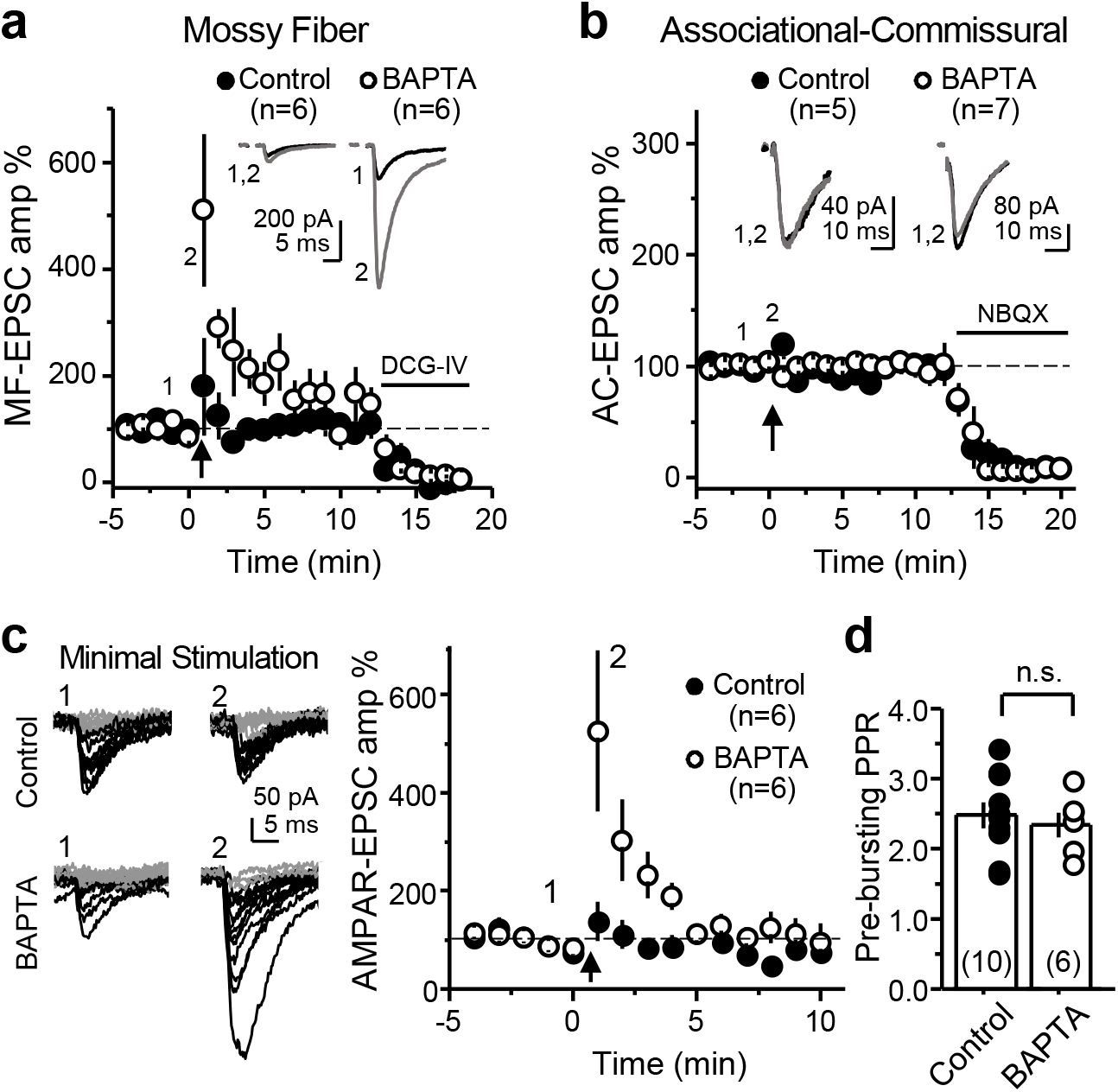
High postsynaptic Ca^2+^ buffering selectively enhances post-tetanic potentiation at the MF-CA3 synapse. (**a**) Summary data showing the effect of loading the patch pipette with 10 mM BAPTA on PTP of AMPAR-mediated MF-responses compared to control condition (0.1 mM EGTA) (Control: 125 ± 52% of baseline; n=6; BAPTA: 347 ± 84%; n=6; Control vs. BAPTA: p<0.05). Experiments were performed at 32 °C, Vh = −60 mV, with only a low concentration of the AMPAR selective antagonist GYKI-53655 (1 μM), and the bursting paradigm was performed in current-clamp mode. AMPAR-EPSCs were evoked with bulk stimulation in the GC layer. At the end of the experiments, DCG-IV (1 μM) was added to the bath to verify that the synaptic responses were mediated by MF activation. PTP is quantified as the average of the EPSCs during the first three minutes post-induction vs the three minutes prior induction. Representative traces (*top*) and time-course summary plot (*bottom*). (**b**) Summary data of experiments in which the same bursting paradigm was performed at the neighboring associational-commissural (CA3-CA3) synapse. Note no difference between control and BAPTA-dialyzed cells (Control: 101 ± 6%; n=5; BAPTA: 95 ± 3%; n=7; Control vs. BAPTA: p>0.3). NBQX was used at the end of the experiments to confirm AMPAR-mediated responses. (**c**) PTP assessed by minimal stimulation of MFs while monitoring AMPAR-EPSCs in the absence of drugs in the bath. PTP was induced in current-clamp mode. *Left*, representative traces showing successes (black) and failures (grey). *Right*, summary data showing MF-PTP elicited with minimal stimulation in control and BAPTA-dialyzed cells (Control: 112 ± 27 of baseline; n=6; BAPTA: 354 ± 82; n=6; Control vs. BAPTA: p < 0.05). (**d**) PPR was not affected by intracellular 10 mM BAPTA loading (Control: 2.5 ± 0.18; n=10; BAPTA: 2.3 ± 0.17; n=6; Control vs. BAPTA: p > 0.5). Number of cells are indicated between brackets. Here and in all figures, data are presented as mean ± SEM; representative traces correspond to the time points indicated by numbers on the time-course plots.

Responses were generated by stimulating presynaptic axons and recording from CA3 pyramidal neurons. Kainate receptor (KAR) responses were recorded in the presence of GYKI-53655 (30 μM) or LY 303070 (15 μM), and CGP-55845 (3 μM) in the bath, and MK-801 (2 mM) in the pipette. With the exception of data obtained in **Fig. 1a, Extended Fig. 5c** and **Fig. 6e–h**, α-amino-3-hydroxy-5-methyl-4-isoxazole propionic acid receptor (AMPAR)–mediated responses were also generated with MK-801 in the pipette, and GYKI-53655 (1 μM) in the bath (Kwon and Castillo, 2008a). N-methyl-D-aspartate receptor (NMDAR) responses were recorded with NBQX (10 μM) in the bath. All experiments except those in **Fig. 1a,c** and **Fig. 6e–h** contained picrotoxin (100 μM).

KAR and NMDAR-mediated EPSCs were evoked by placing a monopolar stimulating pipette with a broken tip (~5-10 μm diameter, filled with ACSF) in the dentate gyrus (DG) cell body layer. For AMPAR–mediated responses, the tip was left unbroken (~1 μm) to minimize the number of MFs activated. For minimal stimulation experiments, a theta-glass stimulating pipette was placed in stratum lucidum 100 μm apart from the recorded CA3 pyramidal cell. Intensity was increased until a success/failure pattern of AMPAR-EPSC responses was observed. MF AMPAR-EPSCs were only accepted for analysis if the following criteria were met: robust PPF (at least 2 fold), DCG-IV sensitivity > than 85%, fast rise time (10-90%) was < 1.2 ms, response onset was < 5.0 ms, and these values did not significantly change after the bursting. These criteria were based on those established by previous studies of MF-CA3 transmission (Jonas et al., 1993). To record KAR- and AMPAR-EPSCs, cells were voltage-clamped at −60 mV. For NMDAR-EPSCs, cells were held at −50 mV. Unless otherwise stated, baseline NMDAR- and KAR-EPSCs were obtained by delivering two stimuli separated by 5 ms in order to evoke a measurable response (Weisskopf and Nicoll, 1995). Stimulus intensity for KAR/NMDAR- and AMPAR-mediated responses was approximately 100 and 10 μA, respectively. Stimulus intensities did not differ, on average, between control and BAPTA-dialyzed cells in any given condition. Stimulus duration was 100-200 μs. CA3 neurons were always dialyzed for ≥ 15 min, while stimulating at 0.1 Hz, before delivering any plasticity-inducing stimulation. For synaptically-evoked action potentials (**Fig. 6e–h**), no drugs were bath applied and CA3 pyramidal cells were held in current-clamp mode before, during and after the induction. Resting potential was kept between −70 and −75 mV. Baseline and post-induction spiking probability were measured as the average of number of spikes per burst normalized to the number of pulses per burst. (i.e. 3 pulses at 25 Hz).

The baseline stimulation frequency for all experiments was 0.1 Hz, except for frequency facilitation (5 stimuli, 25 Hz), which was delivered at 0.05 Hz. The standard PTP induction protocol consisted of 25 bursts (5 stimuli, 50 Hz) delivered at 2 Hz. PTP was never generated more than once in a given slice. To achieve a similar time course of potentiation in BAPTA-dialyzed cells when the inter-stimulus interval was 40 ms (**Fig. 3c**), or when monitoring MF AMPAR-mediated transmission (**Figs. 1a**, **3b**), it was necessary to use 50 bursts. The bursting protocol was delivered while cells were voltage-clamped at −60 mV, except for one experiment in which the induction was performed in current-clamp (**Fig. 1a,c** and **Fig. 6e–h**). Long-term potentiation (LTP) in CA1 pyramidal neurons was induced by pairing postsynaptic depolarization from −60 mV to 0 mV for 3 min with low-frequency stimulation of Schaffer collaterals (180 pulses, 2 Hz).

In all experiments examining MF synaptic transmission, the mGluR2/3 agonist DCG-IV (1-2 μM) was applied at the end of the experiment and data were included only when the response was inhibited by more than 85%. For perforated patch experiments, nystatin was first dissolved into DMSO (10 mg/mL). This was then diluted 250-fold into the intracellular solution to yield 40 μg/mL. Botulinum toxin-B (BoTX) was prepared by making a 0.5 μM stock solution with 1 mg/mL BSA. This was then diluted 100-fold into the final intracellular solution, with 0.5 mM DL-Dithiothreitol (DTT).

### Reagents

MK-801, NBQX, CGP-55845, Nimodipine, AM-251, DCG-IV, and GYKI 53655 were obtained from Tocris-Cookson (Minneapolis, MN, USA). LY 303070 was obtained from ABX advanced biochemical compounds (Radeberg, Germany). BoTX was obtained from List Biological (Cambell, CA, USA). All other chemicals and drugs were purchased from Sigma-Aldrich (St. Louis, MO, USA).

### Data Analysis

Experiments were executed with a MultiClamp 700B amplifier (Molecular Devices, Union City, CA, USA). Data were analyzed online using IgorPro (Wavemetrics, Lake Oswego, OR, USA), and offline with Origin 9.2 (Northampton, MA, USA) and GraphPad Prism (La Jolla, CA, USA). The three minutes before the induction protocol was used as a baseline for statistics. Following the protocol, the first three minutes were used to calculate PTP, and the last three minutes (of a 30 minute period) for the positive control experiment testing LTP at the Schaffer collateral to CA1 pyramidal cell synapse (**Extended Fig. 5c**). Representative responses are averages of 18 traces. All values are shown as mean ± SEM. Unless otherwise stated, Student's t-test was used for statistical significance between two samples, and ANOVA for multiple comparisons. Data that did not display a normal distribution were compared using the non-parametric test Mann-Whitney. All experiments for a given condition were performed in an interleaved fashion – i.e. control experiments were performed for every test experiment.

## RESULTS

### Mossy fiber post-tetanic potentiation is minimal under physiological postsynaptic Ca^2+^ buffering conditions

This study was initiated by the unexpected observation that MF-PTP magnitude was highly dependent on the postsynaptic Ca^2+^ buffering conditions. We induced MF-PTP by activating MFs with a bursting protocol (25 bursts delivered at 2 Hz; 5 stimuli at 50 Hz within a burst) designed to mimic physiological activity patterns of GCs *in vivo* (Henze et al., 2002; Pernia-andrade and Jonas, 2014; Diamantaki et al., 2016; GoodSmith et al., 2017; Senzai and Buzsaki, 2017), while monitoring AMPAR-EPSCs (see Methods) under physiological recording conditions –e.g. no drugs in the bath, near-physiological recording temperature (32°C) and voltage clamping at resting membrane potential (chloride reversal potential). The PTP induction protocol was delivered in current-clamp configuration so that CA3 cells were able to fire freely. To our surprise, we did not observe much potentiation when postsynaptic CA3 pyramidal cells were loaded with 0.1 mM EGTA, a near physiological intracellular Ca^2+^ buffering condition that we refer as “control”, but saw robust MF-PTP with 10 mM BAPTA in the postsynaptic pipette (**Fig. 1a**). In contrast, BAPTA did not increase PTP of associational-commissural (AC) synaptic responses (**Fig. 1b**). These results are consistent with previous studies that reported little potentiation at the AC synapse (Salin et al., 1996), and suggest that the robust PTP unmasked in BAPTA-dialyzed cells is specific to the MF-CA3 synapse. Importantly, the enhancement of MF-PTP under high postsynaptic Ca^2+^ buffering conditions was also observed with minimal stimulation of MFs (Jonas et al., 1993) (**Fig. 1c**), indicating that the PTP enhancement is not an artifact due to strong extracellular stimulation. Lastly, we found that 10 mM BAPTA did not affect the basal paired-pulse ratio (PPR) (i.e. prior to bursting) (**Fig. 1d**), making unlikely that changes in basal Pr could account for the PTP enhancement. These initial observations suggested that MF-PTP, a phenomenon widely believed to be presynaptic in nature, was under the control of a postsynaptic Ca^2+^-dependent process that deserved further investigation.

A major problem when studying MF-CA3 synaptic plasticity is the polysynaptic contamination associated with extracellular stimulation of MFs (Claiborne et al., 1993; Henze et al., 2000; Nicoll and Schmitz, 2005; Kwon and Castillo, 2008a). Repetitive stimulation aggravates this problem as MF-CA3 synapses can be potentiated several-fold by strong frequency facilitation (Regehr et al., 1994; Salin et al., 1996). To rule out the possibility that the PTP was the result of polysynaptic contamination, we blocked AMPA and NMDA receptor-mediated transmission, and monitored kainate receptor-EPSCs (KAR-EPSCs), which are observed at MF-CA3 synapses but not AC-CA3 synapses (Castillo et al., 1997). When CA3 pyramidal neurons were loaded with 10 mM BAPTA the burst stimulation protocol caused a robust PTP of KAR-mediated transmission as compared to control (0.1 mM EGTA) (**Fig. 2a**). To discard that PTP suppression could be due to some unexpected effect of BAPTA, we repeated our experiments with 10 mM EGTA, a slow Ca^2+^ chelator that is widely used at this concentration in voltage-clamp recordings. MF-PTP was equally robust in 10 mM EGTA-loaded cells (**Fig. 2b, Extended Fig. 2**). To ensure fast postsynaptic Ca^2+^ chelation, subsequent experiments compared cells loaded with 0.1 mM EGTA vs. 10 mM BAPTA. Collectively, these findings demonstrate that postsynaptic Ca^2+^ buffering had a striking influence on the magnitude of MF-PTP.

**Figure 2.**
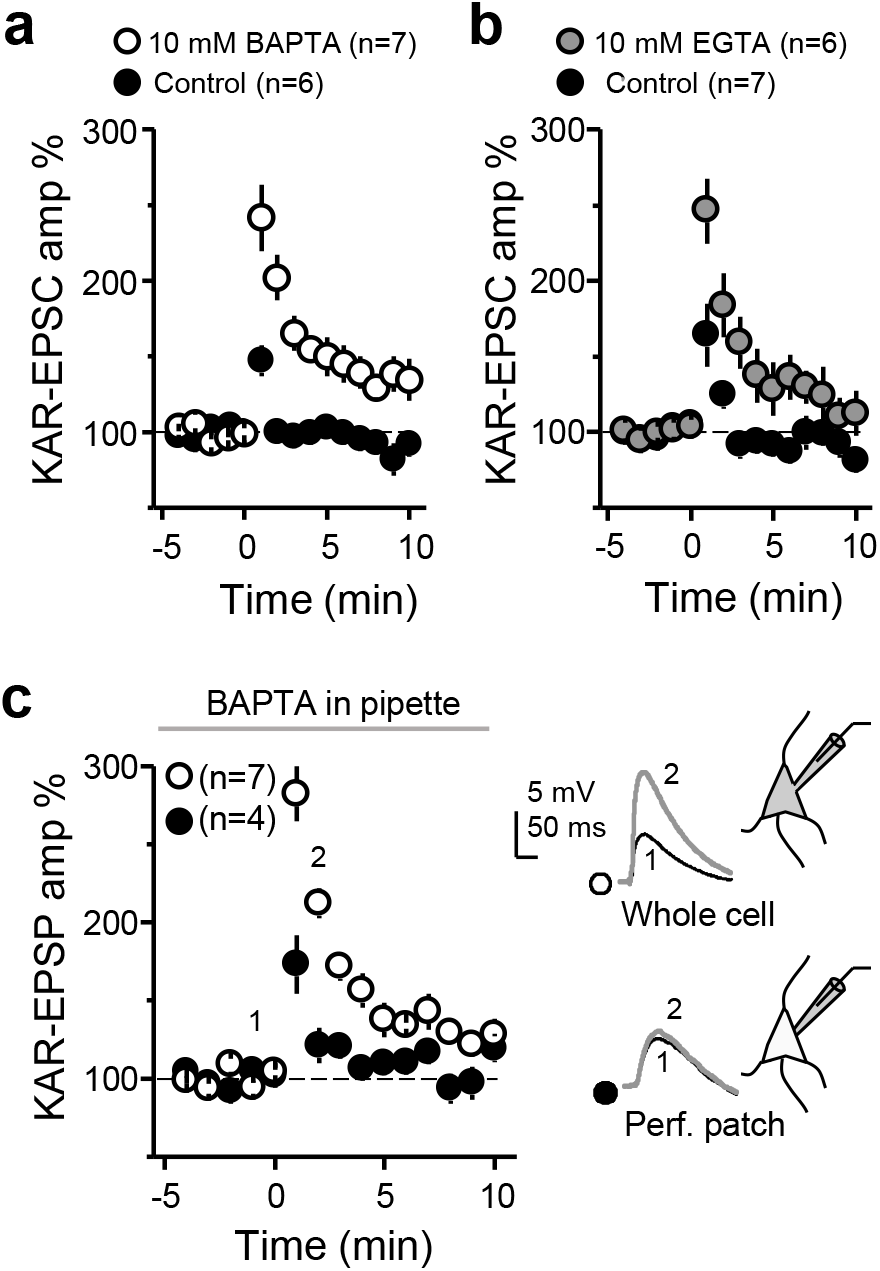
Weak MF-PTP of KAR-mediated transmission under physiological postsynaptic Ca^2+^ buffering recording conditions. (**a**) Summary data showing the effect of 10 mM BAPTA on MF-PTP (Control: 115 ± 6%; n=6; BAPTA: 203 ± 14%; n=7; Control vs. BAPTA: p<0.001). (**b**) Summary data for Control vs 10 mM EGTA (Control: 122 ± 13%, n=7; 10 mM EGTA: 196 ± 17% of baseline, n=6; Control vs 10 mM EGTA p<0.01). (**c**) *Left*, Summary effect when 10 mM BAPTA was included in the recording pipette, in whole-cell vs. perforated-patch configuration (BAPTA perforated patch: 138 ± 11%; n=7; BAPTA whole cell: 226 ± 11%; n=4; BAPTA perforated patch vs. BAPTA whole-cell: p<0.001). *Right*, EPSPs from representative experiments.

One interpretation of the set of observations above is that CA3 pyramidal neurons normally have high Ca^2+^ buffering capacity (i.e. similar to 10 mM EGTA), and replacing these with the 0.1 mM EGTA solution somehow abolished PTP. To directly address this possibility, we monitored KAR-EPSPs with BAPTA in the recording pipette, in perforated patch vs. whole-cell configuration. In perforated patch conditions, in which BAPTA could not diffuse into the cell, only a small amount of PTP was seen, while KAR-EPSPs in whole-cell mode displayed similar striking potentiation observed above when KAR-EPSCs were recorded from BAPTA-loaded cells (**Fig. 2c**, see also **Fig. 2a**). These results indicate that the endogenous Ca^2+^ buffering capacity of CA3 pyramidal neurons was functionally more similar to the 0.1 mM EGTA solution than to the 10 mM BAPTA solution.

We next confirmed the presynaptic nature of MF-PTP in BAPTA conditions. If this PTP was presynaptic, it should be similarly observed when monitoring KAR-EPSCs, NMDAR-EPSCs and AMPAR-EPSCs (unlike the data reported in **Fig. 1a**, AMPAR-EPSCs in these experiments were pharmacologically isolated and recorded under conditions of low network excitability; see Methods). Indeed, when either NMDAR- or AMPAR-mediated EPSCs were monitored, little to no potentiation was observed in control conditions, but the response was markedly increased in BAPTA (**Fig. 3a,b**). Notably, there was no difference when comparing the PTP magnitude of control cells in KA, AMPA, and NMDA receptor groups to each other (ANOVA; F=1.76; p>0.2, DF=15), or when comparing the BAPTA-dialyzed cells in the three receptor groups to each other (ANOVA; F=0.74; p>0.4, DF=17). Thus, the difference in PTP between control and BAPTA persisted regardless of which postsynaptic receptor was pharmacologically isolated. To assess changes in PPR, we monitored KAR-mediated responses. Here, the induction was accompanied by a greater reduction in PPR in BAPTA-dialyzed cells, consistent with an increase in Pr. Importantly, in both control and BAPTA-dialyzed cells, the recovery time course of EPSC amplitude and PPR mirrored each other (**Fig. 3c**), suggestive of a causal relationship between presynaptic Pr and the postsynaptic response magnitude. Together, these independent lines of evidence confirm a presynaptic locus of MF-PTP in BAPTA-dialyzed cells. Again, as for AMPAR-EPSCs (**Fig. 1d**), no significant difference was noted in the degree of basal PPR (i.e. before bursting) between control and BAPTA-dialyzed cells (Control: 2.4 ± 0.2; n=25; BAPTA 2.7 ± 0.2; n=27; Control vs. BAPTA p>0.2; data not shown), supporting the notion that postsynaptic BAPTA loading did not affect basal Pr. The most parsimonious explanation for a robust presynaptic potentiation being unmasked by preventing a rise in postsynaptic Ca^2+^ is a retrograde signal that suppresses potentiation in normal conditions. This interpretation is consistent with the fact that most forms of retrograde signaling so far described require a rise in postsynaptic Ca^2+^ concentration (Fitzsimonds and Poo, 1998; Regehr et al., 2009).

**Figure 3.**
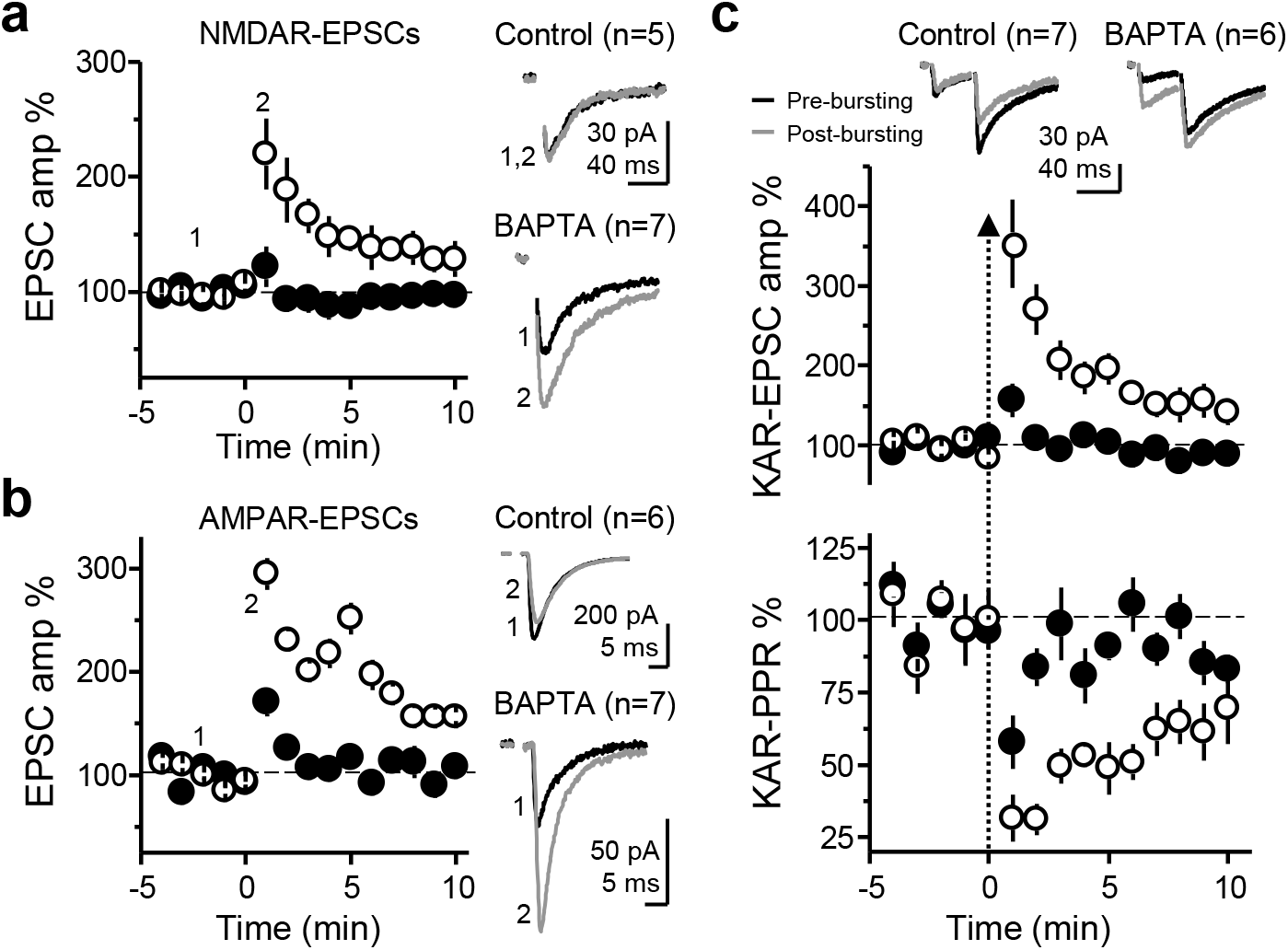
Enhanced MF-PTP by high postsynaptic Ca^2+^ buffering capacity is presynaptically expressed. (**a**) Summary data showing the effect of 25 bursts on NMDAR-mediated EPSCs (V_H=_ −50 mV), when CA3 neurons were dialyzed with control vs BAPTA intracellular solution (control: 102 ± 11% of baseline; n=5; BAPTA: 191 ± 23%; n=7; control vs. BAPTA: p<0.05). (**b**) Summary plot of the AMPAR-mediated MF response after the bursting paradigm, in control vs. BAPTA conditions (control: 135 ± 17%; n=6; BAPTA: 223 ± 16%; n=7; control vs. BAPTA: p<0.01). (**c**) KAR PPR was monitored by delivering two pulses (40 ms inter-stimulus interval). *Top*, Averaged traces from representative experiments, before and after bursting, and summary plot of the first EPSC amplitude in control vs. BAPTA solutions. *Bottom,* Summary of PPR time course as normalized to the three minutes before the bursting paradigm. Same cells as in top panel (Control: 80 ± 8%; n=7; BAPTA: 37 ± 5%; n=6; control vs. BAPTA: p<0.01).

### Source of postsynaptic Ca^2+^ rise involved in PTP suppression

We next sought to assess the source or sources of the postsynaptic Ca^2+^ rise involved in the suppression of MF-PTP. Ca^2+^ influx via NMDARs, or Ca^2+^-permeable AMPARs or KARs, was ruled out as solely sufficient for retrograde suppression, given that pharmacological blockade of these receptors did not unmask PTP (**Figs. 2, 3a**). Several voltage-gated Ca^2+^ channels (VGCCs) exist in the thorny excrescence of CA3 pyramidal neurons (Kapur et al., 2001; Reid et al., 2001). Since the thorny excrescences in our experiments could have been poorly voltage-clamped (at −60 mV) during the induction protocol, one or more of these channels may have contributed to the rise in postsynaptic Ca^2+^. To address this possibility, we clamped CA3 neurons at −80 mV, a voltage at which all VGCCs should be closed. L-type channels mediate a Ca^2+^ rise induced by MF burs stimulation (Kapur et al., 2001). We therefore also added nifedipine (25 μM) to the bath solution to ensure L-type VGCCs were blocked in this experiment. If L-type channels were in fact the source of the Ca^2+^ required for the putative retrograde suppression, we would expect that blocking those channels would mimic the effect of BAPTA, and that both control and BAPTA-dialyzed cells exhibit robust PTP. However, under these recording conditions, a large difference between control and BAPTA-dialyzed cells remained (**Fig. 4a**), suggesting that L-type channels do not contribute significantly to the regulation of MF-PTP. We next examined the potential role of R-type and T-type VGCCs by adding 100 μM Ni^2+^ while blocking L-type VGCCs with 10 μM nimodipine. Under these recording conditions, and likely due to the blockade of presynaptic R-type channels, as previously reported (Breustedt et al., 2003; Dietrich et al., 2003), MF-PTP was dampened, but the difference between low and high postsynaptic Ca^2+^ buffering conditions remained (**Fig. 4b**). It is therefore unlikely that R-type and T-type VGCCs in CA3 pyramidal neurons contribute significantly to the regulation of MF-PTP under physiological intracellular Ca^2+^ buffer conditions.

**Figure 4.**
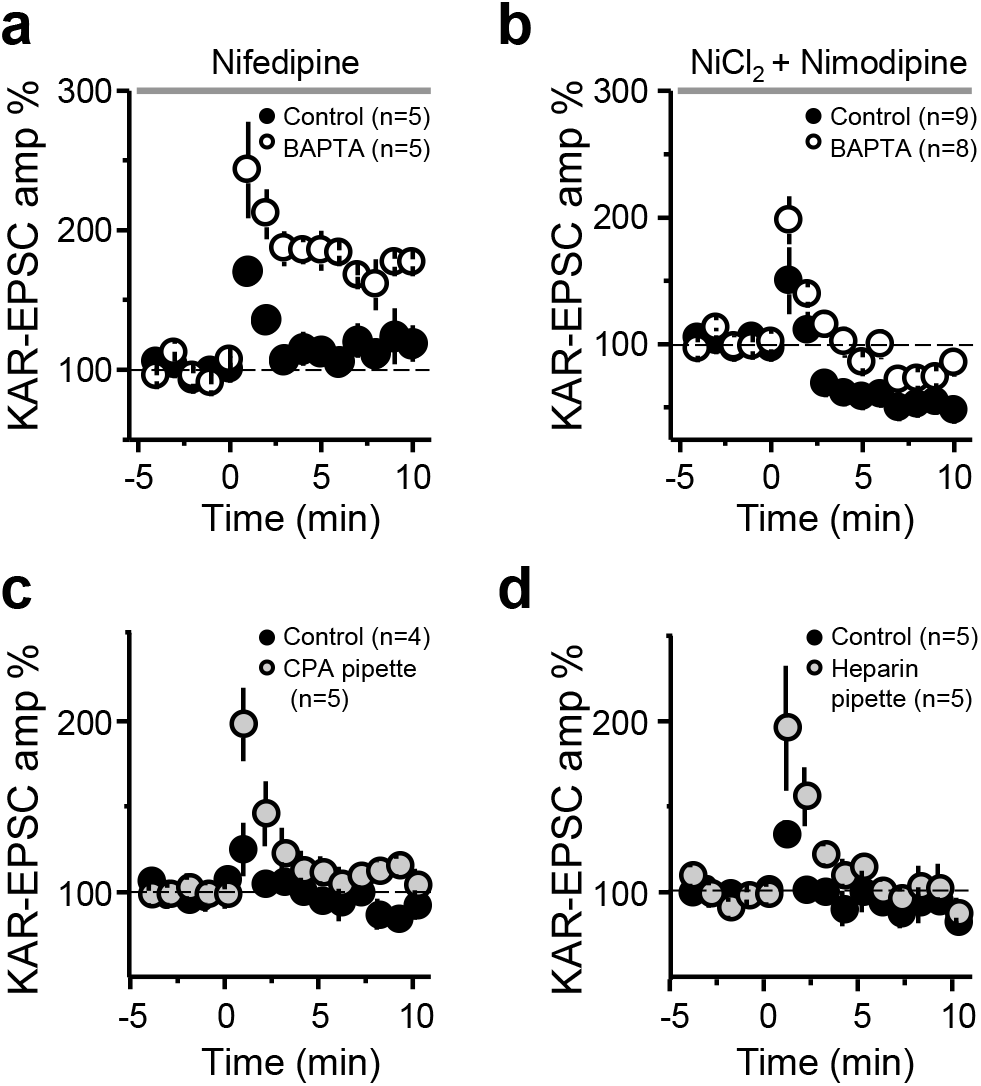
Suppression of MF-PTP depends on internal Ca^2+^ stores. (**a**) Summary effect of voltage-clamping cells at −80 mV with 25 μM nifedipine in the bath in control and BAPTA-dialyzed cells (Control: 138 ± 3%; n=5; BAPTA: 209 ± 20%; n=5; control vs. BAPTA: p<0.01). (**b**) Summary effect of adding 100 μM Ni^2+^ and 10 μM nimodipine to the bath (Control: 110 ± 14; n=9; BAPTA: 151 ± 9; n=8; control vs. BAPTA: p < 0.05). (**c**) Summary effect of including cyclopiazonic acid (CPA) in the patch pipette (Control: 112 ± 6%; n=4; CPA: 156 ± 16%; n=5; Control vs CPA: p<0.05). (**d**) Summary effect of including heparin in the recording pipette to block IP3Rs (Control: 112 ± 2%; n=5; heparin: 157 ± 17%; n=5; Control vs heparin: p<0.05).

To investigate the potential role of intracellular Ca^2+^ stores, we included cyclopiazonic acid (CPA) (30 μM) in the patch pipette to deplete Ca^2+^ from the endoplasmic reticulum. CPA led to an increase in MF-PTP relative to control cells (**Fig. 4c**), suggesting that the rise in Ca^2+^ required by the retrograde signal is mediated, at least partially, by intracellular stores. CPA could have diffused from the recorded cell to the presynaptic terminal, reduced Pr, and thereby increased the magnitude of PTP. To address this possibility, we delivered a synaptic burst (5 pulses, 25 Hz) to cells loaded with control solution, and those loaded with CPA. There was no difference between the ratio of the fifth/first KAR-EPSC amplitude (control: 7.5 ± 1.7; n=5; CPA: 6.9 ± 1.2; n=5; p>0.5; data not shown), suggesting no difference in Pr. Thus, the increase in PTP seen in CPA-loaded cells was likely due to depletion of postsynaptic Ca^2+^ stores. Lastly, release of Ca^2+^ from internal stores can be triggered by activation of inositol 1,4,5-trisphosphate receptors (IP3Rs)(Verkhratsky, 2005), a signaling pathway that has been shown to underlie Ca^2+^ rises in CA3 pyramidal neurons (Kapur et al., 2001) and postsynaptic plasticity (Kwon and Castillo, 2008b) at the MF-CA3 synapse. To examine whether IP3Rs played a role in the suppression of MF-PTP, we included heparin (2.5 mg/mL) in the patch pipette. With IP3Rs blocked, PTP was increased to a similar level as when cells were loaded with CPA (**Fig. 4d**). Together, these results suggest that IP3R-triggered release from internal Ca^2+^ stores contributes to the suppressive effect on PTP.

### Assessing the mechanism underlying MF-PTP suppression

One way that the release of Ca^2+^ from internal stores can be triggered is via activation of group I mGluRs (i.e. mGluR1 and mGluR5 subtypes). These G-protein-coupled receptors (GPCRs) are likely activated during our PTP induction protocol and have been shown to mobilize Ca^2+^ stores at the MF-CA3 synapse (Kapur et al., 2001). However, with the mGluR5 antagonist MPEP (4 μM) and the mGluR1 antagonist CPCCOEt (100 μM) in the bath solution, a large difference remained between the potentiation observed in control and BAPTA-dialyzed cells (**Fig. 5a**), whereas in separate experiments we found these antagonists greatly reduced the inward current induced by the group I mGluR agonist DHPG (**Extended Fig. 5a**). To test for the potential involvement of other GPCRs in mobilizing Ca^2+^ from internal stores (e.g. by activating IP3Rs), we used GDP-βS (1 mM), a nonhydrolyzable GDP analog that interferes with G-protein signaling. Including GDP-βS in the recording pipette solution had no apparent effect on the retrograde suppression as we observed significantly less MF-PTP in control compared to BAPTA-dialyzed cells (**Fig. 5b**). As positive control, we found that intracellularly loaded GDP-βS in CA3 pyramidal neurons abolished the outward current induced by the GABA_B_R agonist baclofen (50 μM) (**Extended Fig. 5b**). Altogether, these results suggest that mobilization of the putative retrograde signal that suppresses MF-PTP requires IP3R-mediated release of Ca^2+^ from internal stores, but is independent from the activation of Group I mGluRs or other GPCR-dependent signaling, although we cannot rule out the possibility that G protein-independent signaling could be involved (Gerber et al., 2007).

Dendrites on postsynaptic neurons have been shown to release retrograde messengers consisting of lipids, gases, peptides, growth factors, and conventional neurotransmitters (Regehr et al., 2009), some of which are released by SNARE-dependent exocytosis. To test whether the putative retrograde signaling suppressing MF-PTP involves vesicular release, we used botulinum toxin-B (BoTX), which cleaves synaptobrevin-2, thus eliminating SNARE-dependent exocytosis (Schiavo et al., 2000; Montal, 2010). Adding BoTX (5 nM) to the intracellular solution did not enhance MF-PTP (**Fig. 5c**). In separate, interleaved experiments that served as a positive control, and as previously reported (Lledo et al., 1998), we found that loading BoTx into CA1 pyramidal neurons blocked LTP of AMPAR-mediated transmission (see Methods) (**Extended Fig. 5c**). These findings argue against SNARE-dependent exocytosis in mediating the putative retrograde signal involved in MF-PTP suppression.

We next examined whether lipids mediate MF-PTP suppression. For instance, endocannabinoids, perhaps the most characterized retrograde signals in the brain (Kano et al., 2009; Castillo et al., 2012), suppress PTP at the parallel fiber-Purkinje cell synapse in the cerebellum by activating presynaptic type 1 cannabinoid receptors (Beierlein et al., 2007). However, these receptors are not expressed at the MF-CA3 synapse in mature animals (Marsicano and Lutz, 1999; Katona et al., 2006; Hofmann et al., 2008; Caiati et al., 2012). To test whether a different lipid signal acting as a retrograde signal, such as arachidonic acid (AA) (Carta et al., 2014), or the AA metabolite 12-(S)-HPETE (Feinmark et al., 2003), could suppress MF-PTP, we added a cocktail of inhibitors to the patch pipette solution in order to inhibit AA and other components of lipid synthesis in the postsynaptic neuron. We included eicosatetraynoic acid (ETYA; 100 μM) and indomethacin (10 μM) to inhibit lipoxygenases and cyclooxygenases (COX 1 and 2), enzymes that catalyze the metabolism of eicosanoids and prostanoids, respectively. We also added RHC-80267 (50 μM) to inhibit diacylglycerol (DAG) lipase. With this combination of lipid inhibitors in the pipette, we continued to see robust MF-PTP in BAPTA-dialyzed cells, but none when the cocktail was included in control cells (**Fig. 5d**). Our results suggest that the putative retrograde signal that suppressed PTP at the MF-CA3 synapse does not depend on these lipid-derived pathways.

Lastly, we explored potential ways by which glutamate release was suppressed during PTP. Presynaptic type 1 adenosine receptors (A1Rs) can tonically inhibit glutamate release at this synapse (Moore et al., 2003) (but see Kukley et al., 2005). To test whether these receptors mediate MF-PTP suppression, we used the A1R-selective antagonist DPCPX (200 nM). DPCPX did not alter the robust difference in PTP observed in control vs. BAPTA-dialyzed cells (**Fig. 5e**), but significantly increased the amplitude of MF KAR-EPSCs (**Extended Fig. 5d**), indicating that DPCPX was active and therefore, A1Rs were not underlying the suppression of MF-PTP. Glutamate release from MFs is also blocked by the activation of presynaptic group II/III mGluRs (Kamiya et al., 1996). While our results using BoTX make it unlikely that these receptors were targeted by glutamate released from the postsynaptic cell, glutamate could have been generated from other sources (e.g. glia). However, in the presence of the group II and III mGluR antagonists LY 341495 (1 μM) and MSOP (200 μM), respectively, the large difference between control and BAPTA remained (**Fig. 5f**). These antagonists almost completely reverse the DCG-IV-mediated suppression of MF transmission (**Extended Fig. 5e**). It is therefore unlikely that activation of presynaptic mGluR2/3 underlies the suppression of MF-PTP.

**Figure 5.**
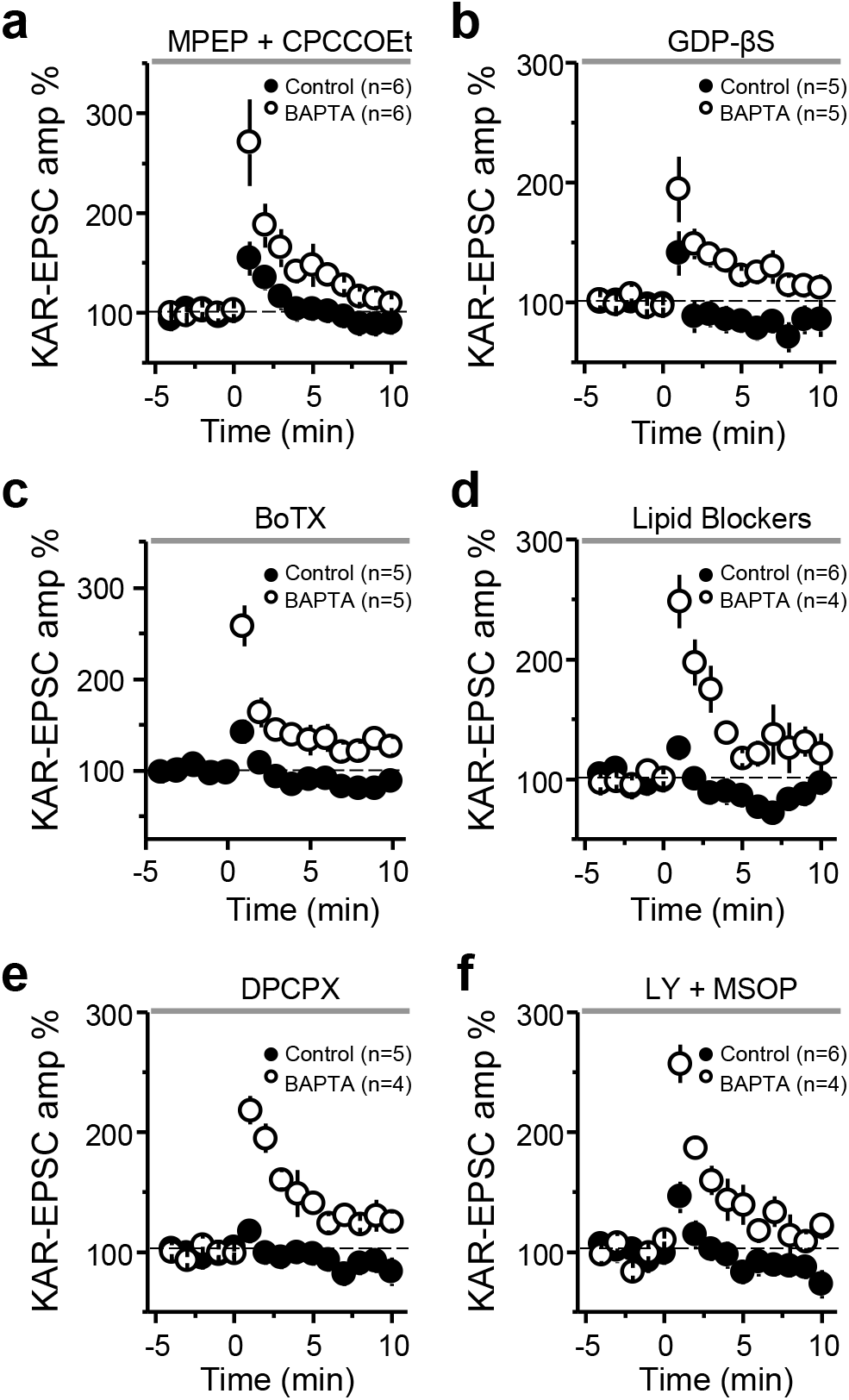
Assessing the mechanism underlying MF-PTP suppression. (**a**) Summary of data of MF-PTP under control (0.1 mM EGTA) and high intracellular buffering conditions (10 mM BAPTA) in the presence of the Group I mGluR antagonists MPEP and CPCCOEt (Control: 122 ± 6%; n=6; BAPTA: 208 ± 28%; n=6; control vs. BAPTA: p<0.05). (**b**) Summary data of experiments in which control and BAPTA cells were dialyzed with GDP-βS (Control-GDP-βS: 106 ± 13%; n=5; BAPTA-GDP-βS: 161 ± 17%; n=5; control-GDP-βS vs. BAPTA-GDP-βS: p<0.05). (**c**) Cells dialyzed with botulinum toxin B (BoTX) did not exhibit any more PTP than interleaved controls without BoTX (BoTX included in control solution: 116 ± 2%; n=5; BAPTA without BoTX: 189 ± 16%; n=5; control-BoTX vs. BAPTA: p<0.01). (**d**) Summary data when a cocktail of lipid blockers (ETYA, indomethacin, and RHC-80267) was included in the intracellular recording solution (Control: 105 ± 6% of baseline; n=6; BAPTA: 220 ± 21%; n=4; control vs. BAPTA: p<0.001). (**e**) Summary data showing no PTP in control cells after bursting paradigm in the presence of the adenosine 1 receptor antagonist DPCPX, whereas robust PTP observed in BAPTA-dialyzed cells (Control: 104 ± 3%; n=5; BAPTA: 191 ± 7%; n=4; control vs. BAPTA: p<0.001). (**f**) Summary data showing the effect of bursting paradigm in control and BAPTA-dialyzed cells in the presence of the Group II and III mGluR antagonists LY-341495 and MSOP (Control: 121 ± 10%; n=6; BAPTA: 201 ± 10%; n=4; control vs. BAPTA: p<0.001).

### Strong presynaptic activity is required for retrograde suppression of glutamate release

The types of activity under which retrograde suppression of transmitter release manifests could have important implications for the CA3 network. We found that burst-induced facilitation, measured by the ratio of the fifth KAR-EPSC amplitude to that of the first (P5/P1) in a single burst (five pulses, 25 Hz), was not significantly different in control vs BAPTA conditions (**Fig. 6a**). We also examined the effect of postsynaptic Ca^2+^ buffering on low-frequency facilitation (LFF), whereby single KAR-EPSCs were evoked, first during 0.1 Hz basal stimulation, and after switching to 1 Hz. No difference was observed between conditions (**Fig. 6b**). Thus, for these modest increases in activity, postsynaptic Ca^2+^ buffering seemed to have no impact on presynaptic transmitter release. Moreover, because frequency facilitation is highly dependent on the starting Pr (Zucker and Regehr, 2002; Nicoll and Schmitz, 2005; Regehr, 2012), these results argue against a tonic suppression of neurotransmitter release under high postsynaptic Ca^2+^ buffering conditions.

**Figure 6.**
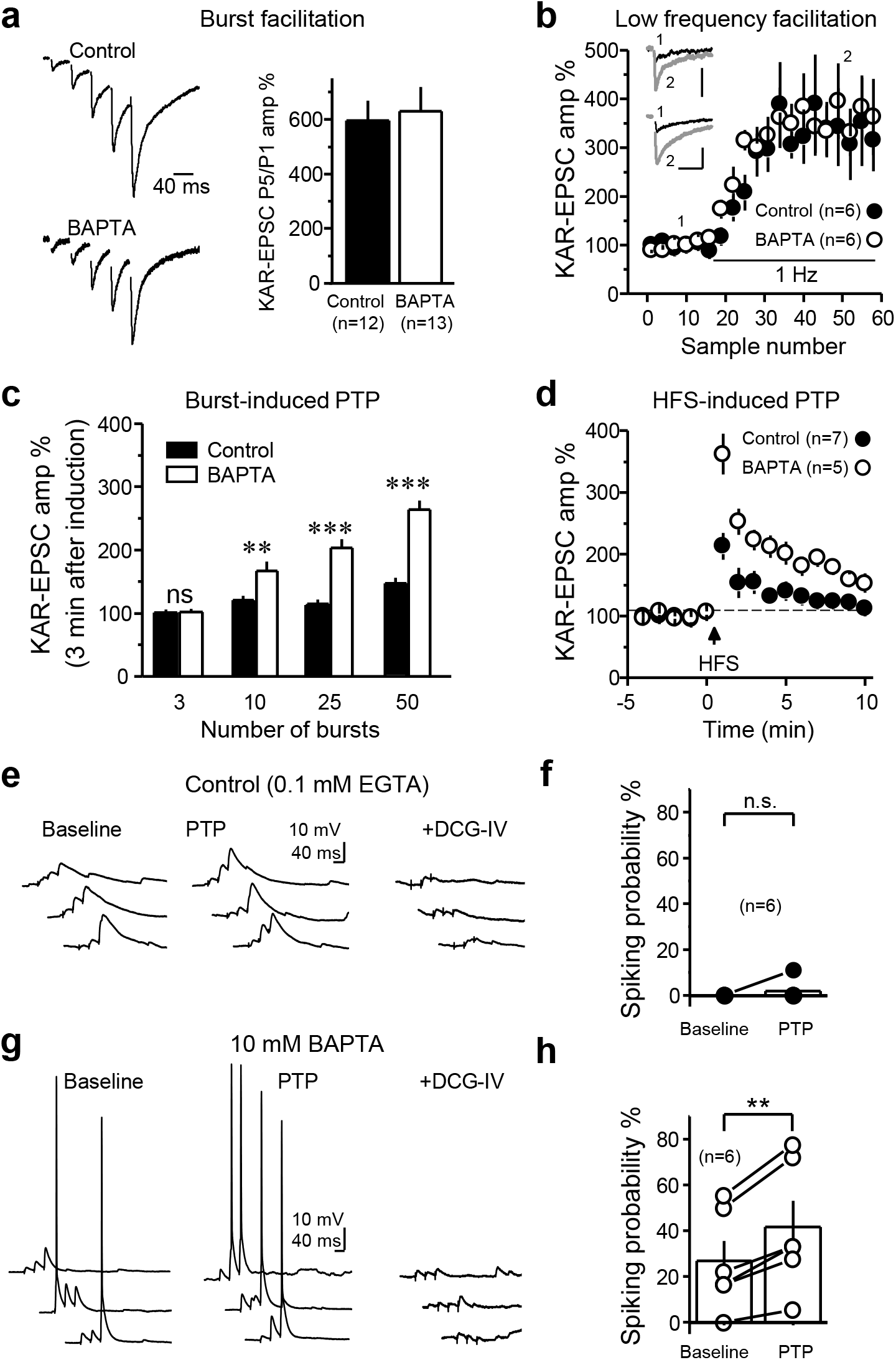
MF-PTP suppression under different patterns of activity. (**a**) Five pulses were delivered to MFs for control and BAPTA-dialyzed cells, and the ratio of the fifth KAR-EPSC to the first was taken (P5/P1 control: 6.0 ± 0.7; n=12; P5/P1 BAPTA: 6.5 ± 0.9; n=13; control vs. BAPTA: p>0.5). Traces normalized to the amplitude of first EPSC. (**b**) Summary data showing effect of switching basal stimulation frequency from 0.1 to 1.0 Hz in control vs. BAPTA-dialyzed cells (LFF control: 341 ± 73% of baseline; n=6; LFF BAPTA: 358 ± 59%; n=6; control vs. BAPTA: p>0.5). Insets: superimposed traces taken at 0.1 and 1.0 Hz. Calibration bars: 40 pA and 40 ms. (**c**) Summary data showing the suppressive effect on MF-PTP induced by different number of bursts. Three bursts, control: 101 ± 5% of baseline; n=7; BAPTA: 102 ± 4%; n=5 (control vs. BAPTA: p>0.5). Ten bursts, control: 121 ± 7% of baseline; n=8; BAPTA: 167 ± 14%; n=7 (control vs. BAPTA: p<0.01). Twenty-five bursts: control: 115 ± 6% of baseline; n=7; BAPTA: 203 ± 14%; n=7 (control vs. BAPTA: p<0.001). Fifty bursts: control: 147 ± 10% of baseline; n=8; BAPTA: 264 ± 15%; n=6 (control vs. BAPTA: p<0.001). (**d**) Effect of high frequency stimulation (HFS; 100 pulses at 100 Hz, x3) on MF KAR-EPSCs, with or without BAPTA in the patch pipette (Control: 181 ± 20% of baseline; n=7; BAPTA: 282 ± 24%; n=5; control vs. BAPTA: p<0.01). (**e-h**) Effects of MF-PTP on CA3 pyramidal neuron firing induced by MF bursting stimulation. No drugs were added to the bath .Control cells loaded with 0.1 mM EGTA (e,f) showed little-to-no action potentials before and after PTP (Baseline: 0 ± 0%; n=6; PTP: 2 ± 2% ; n=6; baseline vs. PTP: p > 0.4, Mann-Whitney), whereas in cells loaded with 10 mM BAPTA (g,h) PTP enhanced the probability of firing action potentials (baseline: 27 ± 9%; n=6; PTP: 42 ± 11%; n=6; baseline vs. PTP: p < 0.005, paired t-test).

We next delivered multiple bursts in order to determine how strong the bursting paradigm must be before retrograde suppression of MF-PTP is observed. To this end, we increased the number of bursts while maintaining both the frequency within a burst (50 Hz), as well as between bursts (2 Hz). After three bursts, synaptic responses were similar in both Ca^2+^ buffering conditions (i.e. control and BAPTA), but a difference emerged after 10 bursts (**Fig. 6c**), and a larger difference was also seen after increasing the number of bursts to 50. In BAPTA-dialyzed cells, there was a difference between 3 vs.10 (ANOVA, F=21.9; p<0.05; DF=10), 3 vs. 25 (p<0.001, DF=10), 3 vs. 50 (p<0.001, DF=9), 10 vs. 50 (p<0.001, DF=11), and 25 vs. 50 bursts (p<0.05, DF=11). Together these data not only uncover the magnitude of MF-PTP in the absence of a retrograde suppressive signal, but also show that in our BAPTA conditions, a longer bursting paradigm induces stronger MF-PTP. The threshold observed with 10 bursts is relatively modest, highlighting that this form of regulation could likely manifest *in vivo*.

We next examined whether stronger activation of MFs could overcome the suppressive retrograde signal. To address this possibility we delivered high-frequency stimulation (HFS) consisting of three trains of 100 stimuli (100 Hz within a train; trains separated by 10 seconds). While a sizeable potentiation was elicited in control cells, the magnitude of MF-PTP was significantly larger in conditions of high Ca^2+^ buffering (**Fig. 6d**). Thus, the putative retrograde signal is strong enough to dampen the PTP evoked even by prolonged high-frequency tetanus.

It has been suggested that the MF-CA3 synapse can operate as a conditional detonator (Treves and Rolls, 1992; Urban et al., 2001; Henze et al., 2002), and a recent study demonstrated that MF-PTP could convert MF-CA3 synapses into full detonators (Vyleta et al., 2016). However, recordings in this study –like many other voltage-clamp studies– were performed under high postsynaptic Ca^2+^ buffering conditions, i.e. 10 mM EGTA in the recording pipette. We therefore reassessed the role of PTP in MF detonation using physiological intracellular Ca^2+^ buffering. To this end, we tested whether PTP induction facilitated the ability of a short MF burst (3 stim, 25 Hz) to generate action potentials in the postsynaptic CA3 pyramidal neurons loaded with either 0.1 mM EGTA (control) or 10 mM BAPTA. Eliciting action potentials in BAPTA neurons was easier than in control neurons, which could be due to changes in excitability (Nelson et al., 2003; Roussel et al., 2006). We found that the spike probability 3 min post-PTP was significantly enhanced in BAPTA but not control cells (**Fig. 6e-h**). These results indicate that the contribution of PTP to MF detonation can be overestimated under high postsynaptic Ca^2+^ buffer recording conditions.

## DISCUSSION

We report here that the Ca^2+^ buffering capacity of the postsynaptic neuron can significantly impact presynaptically-expressed PTP at the hippocampal MF-CA3 synapse. Under normal Ca^2+^ buffering capacity, as observed during non-invasive recording conditions (e.g. perforated patch recording) that do not significantly alter the physiological intracellular milieu, we found that a bursting induction protocol designed to mimic *in vivo* activity patterns of GCs triggered a remarkably weak PTP. However, a far greater potentiation was revealed under high postsynaptic Ca^2+^ buffering conditions. Remarkably, increasing the postsynaptic buffer capacity had no significant effect on the basal Pr, arguing against a tonic suppression of neurotransmitter release. The most parsimonious explanation for our findings is the presence of a Ca^2+^-dependent, retrograde signaling mechanism that suppresses PTP. A minimum threshold was required before the phenomenon was observed, above which it operated in a wide range of activity. These results point to a novel, activity-dependent form of negative feedback at the MF-CA3 synapse that may significantly impact DG – CA3 information transfer.

At first glance, our findings contrast starkly with numerous studies that have reported pronounced MF-PTP (for a review see Henze et al., 2000). However, most of these studies elicited MF-PTP with strong repetitive stimulation (e.g. HFS) while monitoring MF transmission with extracellular field recordings. Due to activation of the CA3 network, these experimental conditions not only enable the recruitment of associational-commissural inputs that are commonly interpreted as MF-mediated responses, but also facilitate population spike contamination of (extracellularly recorded) synaptic responses (Henze et al., 2000; Nicoll and Schmitz, 2005). As a result, MF-PTP magnitude can be easily overestimated. Strong MF-PTP was also observed in studies that used more sensitive, single-cell recordings (see for example Zalutsky and Nicoll, 1990; Maccaferri et al., 1998; Vyleta et al., 2016; Vandael et al., 2020), but here again it was typically evoked with a rather non-physiological induction protocol such as HFS. Critically, many such previous studies also loaded postsynaptic neurons with 10 mM EGTA, a standard concentration used in whole-cell studies, both *in vitro* and *in vivo*. Indeed, we also observed striking MF-PTP in those conditions (**Fig. 2b**). Our findings therefore suggest that MF-PTP is tightly controlled when afferent stimulation and postsynaptic Ca^2+^ buffering are set to more physiological levels.

A rise in postsynaptic Ca^2+^ is required for most forms of retrograde signaling (Fitzsimonds and Poo, 1998; Regehr et al., 2009). Consistent with this notion, MF-PTP was suppressed when the postsynaptic cell was loaded with high (milimolar) concentrations of EGTA or BAPTA, or with CPA or heparin, all agents that prevent a rise in intracellular Ca^2+^. To determine the potential source(s) of postsynaptic Ca^2+^ rise involved in the suppression of MF-PTP, we pharmacologically interfered with these sources one by one. We found that IP3R-mediated Ca^2+^ release from intracellular stores contributed to the suppression, but that Group I mGluRs and G-protein-coupled signaling alone were insufficient. Importantly, however, we cannot discard synergism of multiple sources leading to the full rise in postsynaptic Ca^2+^ required for retrograde suppression of MF-PTP, including the contribution of VGCCs in a poorly voltage-clamped postsynaptic compartment (Williams and Mitchell, 2008; Beaulieu-Laroche and Harnett, 2018). Interestingly, the endogenous Ca^2+^ buffering power of CA3 pyramidal neurons is the source of some debate, as some have suggested it is higher than that of CA1 cells (Wang et al., 2004), while others report it is similar (Simons et al., 2009). Further complicating this matter, different endogenous buffers and extrusion mechanisms create short-lived Ca^2+^ nanodomains which can limit the interaction between Ca^2+^ and its substrates (Higley and Sabatini, 2008). Our data show little PTP when the intracellular composition was not perturbed in perforated patch mode, and robust PTP when the cell was dialyzed with a high concentration of BAPTA (**Fig. 2c**). This observation would indicate that, regardless the exact endogenous buffering power of these cells, it is sufficiently low to allow for the postsynaptic neuron to regulate MF-PTP.

Retrograde signaling has been proposed to influence PTP, but typically in the opposite direction to our findings. For example, in the *Aplysia* sensory-motor neuron preparation, injection of BAPTA into the postsynaptic cell depressed PTP (Bao et al., 1997). In the hippocampus, loading CA3 pyramidal neurons with high concentrations of BAPTA (30-50 mM) curtailed PTP and abolished LTP at the MF-CA3 synapse, presumably by blocking the mobilization of a retrograde signal (Yeckel et al., 1999; but see Mellor and Nicoll, 2001). There is also evidence that the receptor tyrosine kinase ephrin B (EphB) and its membrane-bound ligand ephrin B mediate retrograde communication at MF-CA3 synapses, and that interfering with EphB/ephrin B signaling inhibits both PTP and LTP at MF-CA3 synapses (Contractor et al., 2002; Armstrong et al., 2006). Our findings resemble the endocannabinoid-mediated suppression of PTP at the parallel fiber-Purkinje cell synapse via presynaptic type 1 cannabinoid receptors (Beierlein et al., 2007). However, these receptors are not found at MF boutons in the mature brain (Marsicano and Lutz, 1999; Katona et al., 2006; Hofmann et al., 2008; Caiati et al., 2012). While the identity of the putative retrograde signal generated by the postsynaptic neuron remains unidentified, some usual candidates (Regehr et al., 2009) could be discarded. Vesicular, SNARE-dependent exocytosis was ruled out (**Fig. 5c**), indicating that the messenger is likely not a conventional neurotransmitter (e.g. glutamate or GABA). Further evidence against glutamate as a retrograde signaling mediating PTP suppression is the fact that this suppression remains intact after blocking presynaptic group II and III mGluRs (**Fig. 5f**). Our findings also argue against the role of another putative signal, nitric oxide (NO) given that its synthesis typically requires the activation of NMDARs (Christopherson et al., 1999; Sattler et al., 1999), which were blocked in most of our experiments. We cannot discard the possibility that PTP suppression involves retrograde signaling via synaptic adhesion molecules. It is worth noting that identifying endocannabinoids as the retrograde messengers mediating the well characterized depolarization-induced suppression of inhibition (DSI) took a decade of work by several groups (Barinaga, 2001). Future studies will have to determine the identity of the presumably non-conventional retrograde signal and the presynaptic substrate responsible for curtailing PTP (Regehr, 2012).

Two recent studies suggested that MF-PTP occurs *in vivo* (Vandael et al., 2020) and switches MF-CA3 synapses into full detonators (Vyleta et al., 2016). However, our results indicate that the impact of PTP was likely overestimated because recordings in both studies were performed under high, non-physiological Ca^2+^ buffering conditions (e.g. 10 mM EGTA). PTP has also been reported at MF to inhibitory interneuron synapses (Alle et al., 2001; Mori et al., 2007). Given that MFs make 10 times as many contacts onto inhibitory interneurons as onto CA3 pyramidal cells (Acsady et al., 1998), PTP at MF to interneuron synapses, by presumably activating an inhibitory network, could control the activity of GCs (feed-back inhibition) and detonation of CA3 pyramidal neurons (feed-forward inhibition) (Lawrence and McBain, 2003). However, unlike PTP at MF-CA3 synapses, postsynaptic Ca^2+^ buffering had no impact on PTP magnitude at MF to basket cells synapses in the DG, with similarly robust PTP when the interneuron was loaded with 0.1 mM EGTA or 10 mM BAPTA (Alle et al., 2001). Thus, although PTP is observed at distinct MF synapses, the retrograde suppression we report in the present study seems to be unique to the MF-CA3 pyramidal cell synapse.

Relatively strong GC burst activity is needed for engaging suppression of MF-PTP, and the magnitude of this suppression does not seem to be overcome by stronger presynaptic activity (**Fig. 6c**). Thus, PTP suppression may serve as a powerful mechanism of controlling DG-CA3 information transfer following repetitive bursts of GC activity that normally occur during exploratory behaviors (Henze et al., 2002; Pernia-Andrade and Jonas, 2014; Diamantaki et al., 2016; GoodSmith et al., 2017; Senzai and Buzsaki, 2017). Left unchecked, MF-PTP could indirectly lead to runaway activity of the CA3 network and facilitate epileptic activity. Lastly, while the generalizability of PTP suppression remains untested, future studies will be required to determine whether Ca^2+^ indicators widely used *in vitro* and *in vivo* throughout the brain may promote a similar suppression of presynaptic function.

## Acknowledgements

We thank all the Castillo lab members for invaluable discussions. We also thank Professor Peter Jonas for his insightful comments on a recent version of our manuscript.

---

**Extended Figure 2.**
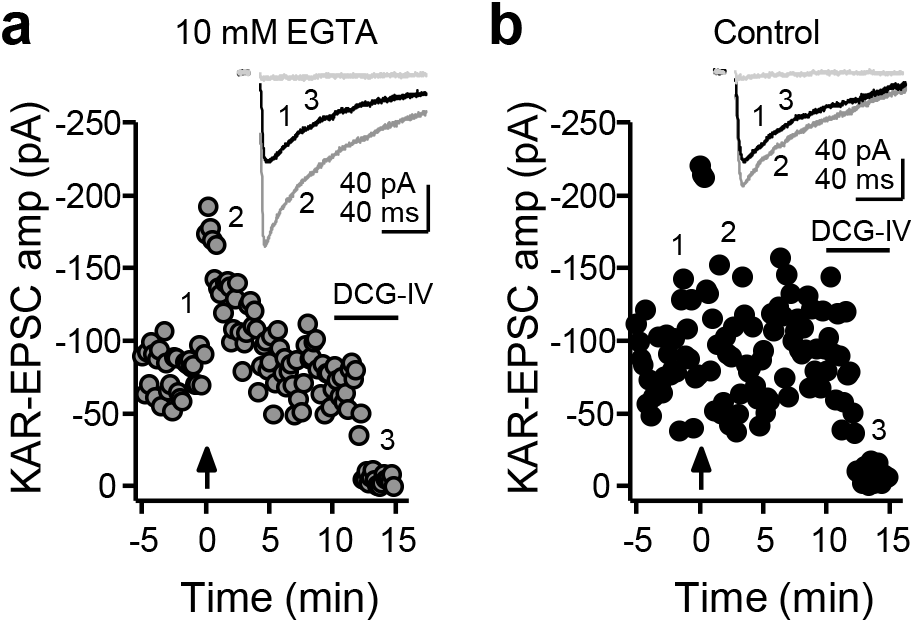
(**a**) Representative experiment in which the recording pipette solution contained 10 mM EGTA. KAR-EPSC traces, which correspond to the numbers in the time course plot below, are shown above. DCG-IV (1 μM) was added at the end of the experiment. (**b**) Representative experiment in which the pipette solution contained 0.1 mM EGTA (Control). Note suppression of PTP relative to (a).

**Extended Figure 5.**
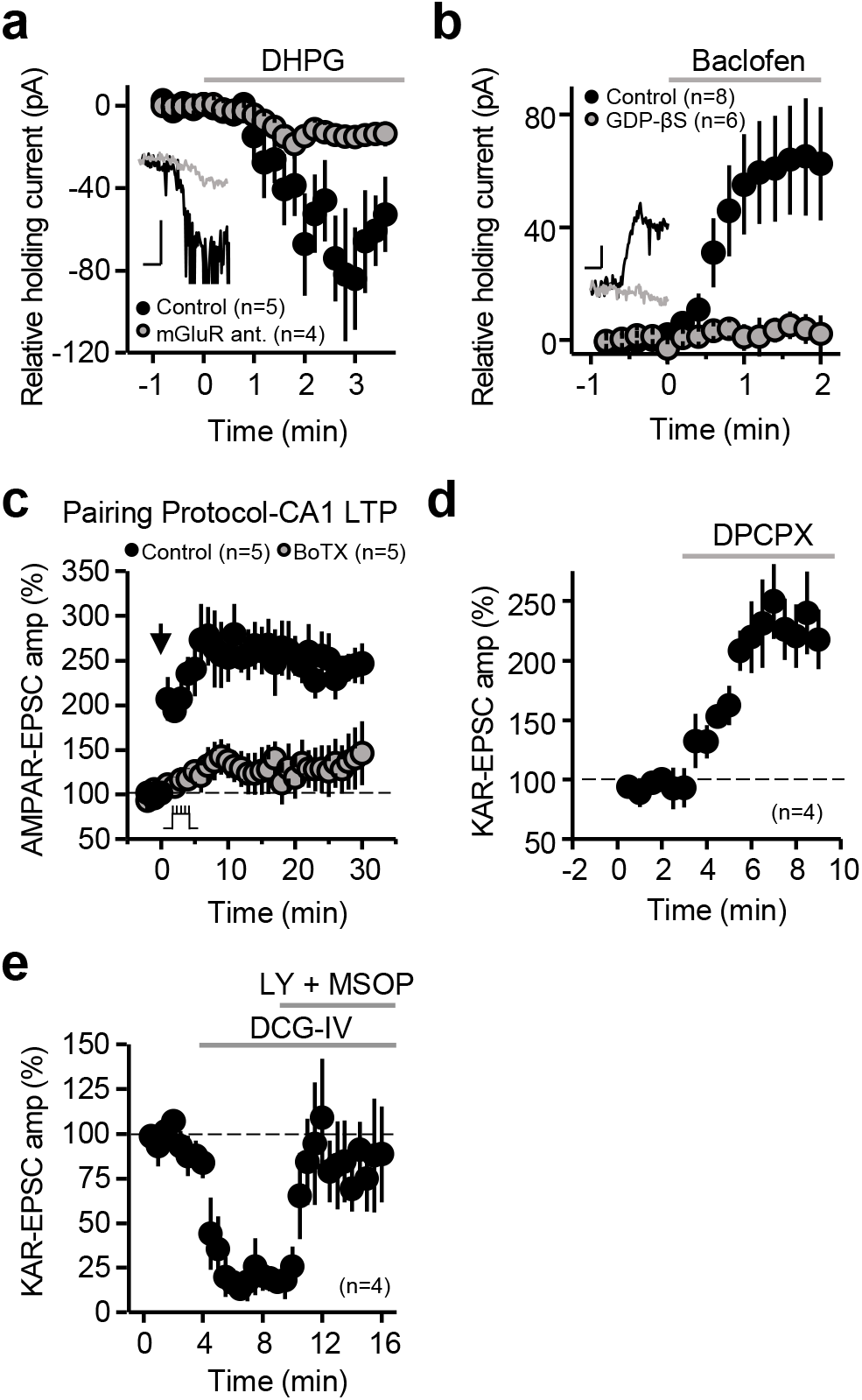
(**a**) Bath application of the group I mGluR agonist DHPG (50 μM) caused an inward current, was significantly reduced in the presence of the group I mGluR antagonists MPEP/CPCCOEt. (Control: 55 ± 11 pA; n=5; MPEP/CPCCOEt: 14 ± 5 pA; n=4; control vs. antagonists: p<0.05). Inset: superimposed traces in control and in the presence of MPEP/CPCCOEt. Calibration bars: 40 pA, 1 min. (**b**) The outward current induced by bath application of the GABA_B_ agonist baclofen was abolished by intracellular loading of GDP-βS (Control: 61 ± 19 pA; n=8; GDP-βS: 3 ± 6 pA; n=6; Control vs. GDP-βS: p<0.05). Inset: superimposed representative traces in control cells (black) and cells loaded with GDP-βS. Calibration bars: 40 pA, 1 min. (**c**) Loading BoTX in CA1 pyramidal neurons blocked LTP of AMPAR-mediated transmission (Control: 240 ± 18% of baseline; n=5; BoTX: 137 ± 30%; n=5; Control vs. BoTX: p<0.05). LTP was generated using a pairing protocol of postsynaptic depolarization (from V_H_ = −60 to 0 mV) for three minutes, and stimulating the Schaffer collaterals with 180 pulses (2 Hz). (**d**) Positive control showing the immediate potentiating effect of 200 nM DPCPX (221 ± 25% of baseline, n=4; p<0.01). (**e**) The suppression of MF-mediated responses by 1 μM DCG-IV bath application was reversed by the antagonists LY341495 (1 μM) and MSOP (200 μM) (90 ± 24% of original baseline; n=4).

